# UVB mutagenesis differs in *NRAS*- and *BRAF*-mutant mouse models of melanoma

**DOI:** 10.1101/778449

**Authors:** Robert L. Bowman, Rebecca C. Hennessey, David A. Tallman, Tirzah J. Weiss, Emma R. Crawford, Brandon M. Murphy, Amy Webb, Souhui Zhang, Krista M. D. La Perle, Craig J. Burd, Ross L. Levine, A. Hunter Shain, Christin E. Burd

## Abstract

*BRAF*-mutant melanomas are more likely than *NRAS*-mutant melanomas to arise in anatomical locations protected from chronic sun damage. We hypothesized that this discrepancy in tumor location is a consequence of the differential sensitivity of *BRAF* and *NRAS*-mutant melanocytes to ultraviolet light (UV)-mediated carcinogenesis. We tested this hypothesis by comparing the mutagenic consequences of a single, narrow band ultraviolet-A (UVA; 320-400nm) or ultraviolet-B (UVB; 280-320nm) exposure in mouse models predisposed to *Braf-* or *Nras*-mutant melanoma. Exposures approximated the amount of UVA or UVB energy contained in ~40 minutes of summer sunlight. Tumor onset was accelerated in all UVB-, but only half of UVA- irradiated mice as compared to unirradiated controls. Melanomas from both mouse models, harbored recurrent mutations affecting the RING domain of MAP3K1 and Actin-binding domain of Filamin A irrespective of UV status. Melanomas from UVB-irradiated, *Braf*-mutant mice averaged twice as many SNVs (1,025 vs. 435) and five times as many dipyrimidine variants (33.3 vs. 5.7) than tumors from similarly irradiated *Nras*-mutant mice. We identified a mutational signature enriched in UVB-accelerated tumors which mirrored COSMIC signatures associated with human skin cancer. Notably, this signature was enriched to a greater extent in *Braf-* than *Nras*-mutant murine melanomas. These data suggest that oncogenic BRAF may enhance UVB carcinogenesis to promote melanoma formation at anatomic sites with low or intermittent sun exposure.

## INTRODUCTION

The most common genetic subtypes of human melanoma, *NRAS-* and *BRAF*-mutant, are enriched in different anatomical locations. *NRAS*-mutant melanomas preferentially localize to chronically sun-damaged (CSD) skin on the head and neck, whereas *BRAF*-mutant melanomas are more common in areas of intermittent sun exposure (Zhang et al. 2016). Despite the association of *NRAS*-mutant tumors with CSD skin, it is reported that UV signature lesions (C>T and CC>TT) are prevalent in a similar proportion of *NRAS-* and *BRAF*-mutant melanomas (Cancer Genome Atlas 2015). These observations led us to speculate that *BRAF*-mutant nevi could be more sensitive to UV mutagenesis and subsequent melanocyte transformation. However, it is difficult to control for differences in lifetime sun exposure among biopsies of human melanomas or nevi.

Genetically engineered mouse models (GEMMs) provide a controlled genetic background in which the genomic and phenotypic effects of UV exposures can be studied. GEMMs encoding a melanocyte-specific *Nras-* or *Braf*-mutation mimic the presense of these mutations in human benign nevi (Roh et al. 2015). Moreover, neonatal or chronic UV treatment accelerates the formation and progression of melanoma in a variety of melanoma GEMMs, consistent with human disease etiology (Mukhopadhyay et al. 2016; Chagani et al. 2017; Hennessey et al. 2017; Perez-Guijarro et al. 2017; Trucco et al. 2019). Genomic analyses of tumors from UV-treated *Nras* or *Braf*-mutant GEMMs have been reported (Viros et al. 2014; Mukhopadhyay et al. 2016; Trucco et al. 2019). However, no study has directly compared the mutational profiles of *Nras-* and *Braf-* driven mouse melanomas exposed to the same UV dosing scheme. Therefore, a complete understanding of how different oncogenic drivers cooperate with environmental mutagens to promote transformation is lacking.

Here, we used a single-dose UV irradiation scheme to characterize the phenotypic and genomic effects of narrow band UVA (360-390 nm) and UVB (280-320 nm) exposures in *Nras-* and *Braf*-mutant mouse models of melanoma. We exposed these animals to a single dose of UVA or UVB, approximating the amount of energy from each band of the UV spectrum in 40 minutes of intense sunlight. Then, we monitored the mice for melanoma development. Tumors from these animals were sequenced to gain insight into the mutational consequences of each UV source in *Nras-* and *Braf*-mutant melanocytes.

## RESULTS

### UV exposure alters NRAS- and BRAF-mutant melanomagenesis

We generated melanocyte-specific, Tyr::CreER(T2)-driven, *Nras (TN)* and *Braf (TB)* mice to model the major genetic subtypes of human melanoma (Hodis et al. 2012) (**Figure 1A-B**). *TN* mice are homozygous for the *LSL-Nras^Q61R^* allele (Burd et al. 2014; Hennessey et al. 2017), whereas *TB* animals carry a heterozygous, conditional *Braf^V637E^* allele *(Braf^CA^;*(Dankort et al.2007). Notably, the *Braf^V637E^* allele is the murine equivalent of human *Braf^V600E^*(Rad et al. 2013). Oncogene expression is driven by the endogenous gene promoter in both models, and is activated by a melanocyte-specific, tamoxifen-inducible Cre recombinase (Tyr::CreER(T2); (Bosenberg et al. 2006)). Therefore, the expression of oncogenic *Nras* or *Braf* in these mice mimics the presence of *NRAS* and *BRAF* mutations in most benign human nevi (Roh et al. 2015). Mice carrying only the *Braf^CA^* allele rarely develop melanoma (Dankort et al. 2009). For this reason, *p16^INK4a^* conditional knockout alleles *(p16^L^;*(Monahan et al. 2010)) were included in both the *TN* and *TB* models. While p16^INK4a^ loss-of-function is an early event observed in >60% of human melanomas, germline mutations affecting *p16^INK4a^* are insufficient to drive the disease in mice or humans (Bishop et al. 2000; Shain et al. 2015b; Hennessey et al. 2017).

**Figure 1.**
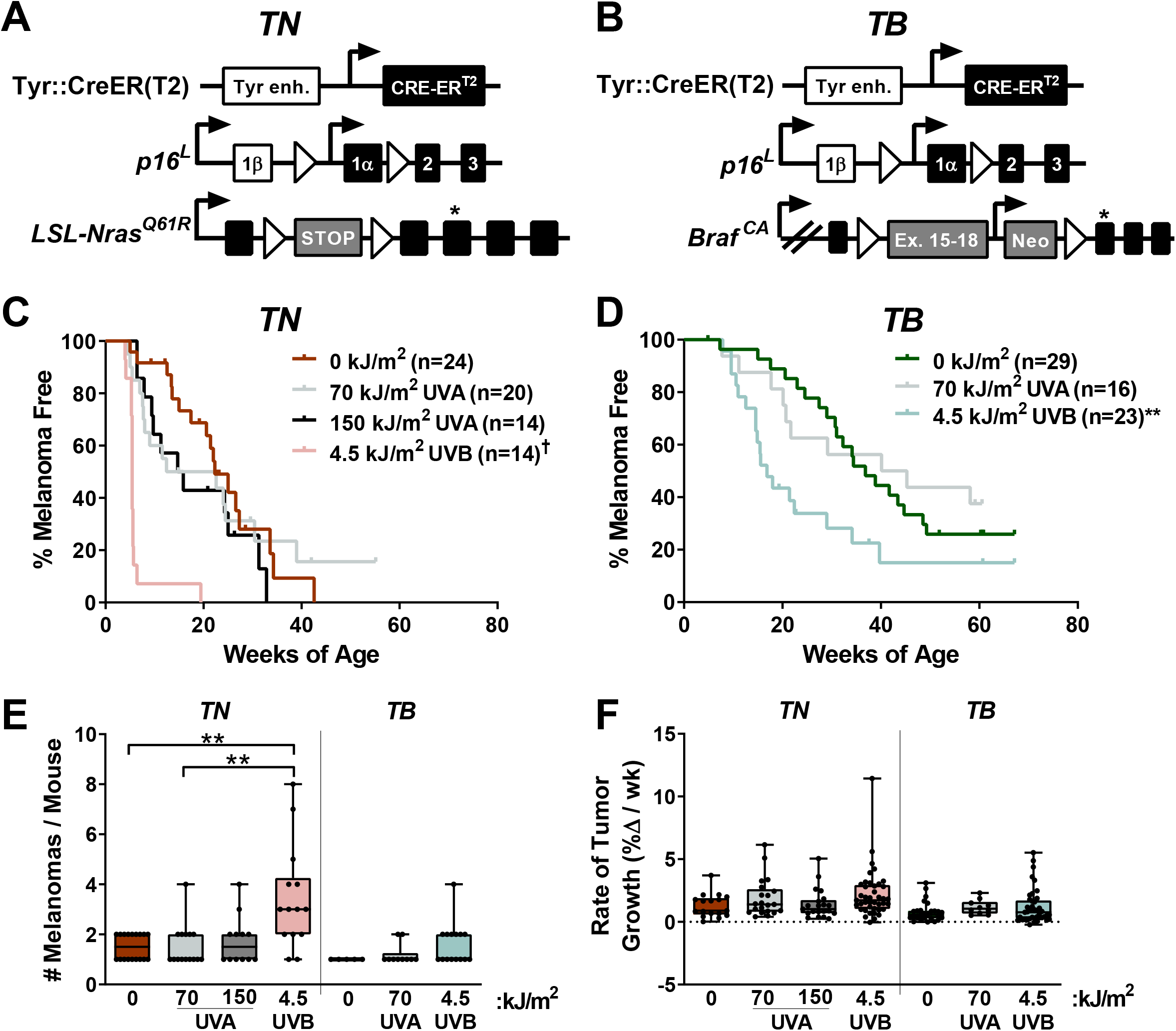
Neonatal UV exposure alters melanoma onset in *TN* and *TB* mice. **A**, *TN* mice are homozygous for a melanocyte-specific, tamoxifen-inducible Cre transgene (Tyr::CreER(T2)), a conditional *p16^INK4a^* knockout allele *(p16^L^),* and a conditional *Nras^Q61R^* knock-in allele *(LSL-Nras^Q61R^).* Open triangles represent LoxP sites. A star indicates the location of the *Nras^Q61R^* mutation. **B**, *TB* mice carry a single, *Braf^V637E^* conditional allele *(LSL-Braf^CA^)* and are homozygous for Tyr::CreER(T2) and *p16^L^.* Note that BRAF^V637E^ is the murine equivalent of human BRAF^V600E^. Open triangles represent LoxP sites and the location of the V637E mutation is indicated by a star. **C & D**, Kaplan-Meier curves depicting the melanoma-free survival of *TN* (C) and *TB* (D) mice treated on postnatal day three with a single dose of ambient light (0 kJ/m^2^), UVA or UVB. † =*p*<0.0001, ** =*p*<0.01 comparing control (0 kJ/m^2^) and UV-irradiated animals of the same genotype (Gehan-Breslow-Wilcoxon). **E**, Total tumor burden of control and UV-irradiated *TN* and *TB* mice at euthanasia. Each circle represents a single mouse. Boxes represent the mean and interquartile range for each group. Whiskers span from the minimum to the maximum value. ** =*p*<0.01 comparing control (0 kJ/m^2^) animals of the same genotype (two-tailed, unpaired t-tests with Welch’s correction). **F**, Average tumor growth rates for UV- and mock-irradiated *TN* and *TB* mice. Each circle represents a single tumor *(TN:* 0 kJ/m^2^ n=17; 70 kJ/m^2^ UVA n=21; 150 kJ/m^2^ UVA n=21; 4.5 kJ/m^2^ UVB n=41; *TB:* 0 kJ/m^2^ n=27; 70 kJ/m^2^ UVA n=10; 4.5 kJ/m^2^, n=41). Data are presented as described in ‘E’.

*TN* and *TB* mice were topically treated with 4-hydroxytamoxifen (4OHT) on postnatal days one and two to induce Cre activity and stimulate recombination of the conditional *p16^INK4a^* knockout and *LSL-Nras^61R^* or *Braf ^CA^* alleles. On postnatal day three, the mice were exposed to a single dose of ambient light (“No UV” or 0 kJ/m^2^), narrow band UVA or narrow band UVB irradiation. The amount of UVB delivered approximated that which is contained in 40 minutes of summer sunlight (4.5 kJ/m^2^), whereas the amount of UVA employed models a short, indoor tanning session (70 or 150 kJ/m^2^; see: **Methods**). These dosing schemes approximate sun exposures of a similar duration, as the UVB to UVA ratio in sunlight is ~1:20, but varies based on season, cloud cover and latitude (Cadet and Douki 2018). Neither dose of UVA or UVB caused erythema or blistering.

The onset of spontaneous melanoma was compared among mice exposed to No UV, UVA or UVB irradiation. Exposure to a single dose of 4.5 kJ/m^2^ UVB dramatically accelerated melanoma onset and decreased overall survival in both the *TN* and *TB* models (**Figures 1C-D**, **Suppl. Figure S1**). Exposure to 70 kJ/m^2^ UVA also enhanced melanoma formation, but only half of the mice developed tumors earlier than the median onset in unirradiated animals (**Figure 1C-D**). Doubling this dose of UVA in the *TN* model did not further facilitate melanoma formation, suggesting that 70 kJ/m^2^ UVA was sufficient to elicit the maximal response achievable with a single exposure (**Figure 1C**). Together, these results reveal the potent ability of narrow band UVB to promote melanoma formation in *TN* and *TB* mice. Furthermore, our findings suggest that UVA could facilitate melanoma onset in some settings, albeit to a much lesser extent than UVB.

We next examined the incidence and growth phenotypes of tumors arising in each of our experimental cohorts. Tumor incidence (# melanomas/mouse) increased in UVB-exposed *TN* mice, but was not significantly altered in *TN* animals treated with UVA or *TB* mice exposed to any form of UV (**Figure 1E**). Tumor distribution and incidence were also similar between male and female *TN* and *TB* mice regardless of exposure, with ~59% of tumors arising on the trunk, ~13% on the head and ~16% on the ears or tail (**data not shown**). Once established, *TN* and *TB* tumors grew at the same rate regardless of prior exposure (**Figure 1F**). Therefore, early tumor onset, rather than more rapid melanoma growth, is responsible for the reduction in overall survival observed in UVB-exposed *TN* and *TB* mice.

We postulated that melanomas arising in UVA- or UVB-exposed mice would exhibit distinct histopathological features. Therefore, we examined hematoxylin and eosin stained tumor sections representative of the rate of onset and body site distribution of melanomas from each cohort. Tumors from both models contained variable percentages of myxoid and spindle cells with comparable degrees of invasion, mitosis and granulocyte infiltration regardless of treatment (**Suppl. Figure S2A-C**, **data not shown**). A paucity of pilosebaceous units and hyperplasia of the overlying epidermis was also observed in UVA, UVB and unexposed mice of both genotypes (**Suppl. Figure S2A&F**). Most tumors from the UVB-treated *TN* cohort contained neoplastic cells with plasmacytoid features that were not prevalent in *TN* melanomas from the UVA and No UV cohorts (6 of 7 vs. 2 of 6 and 0 of 5 tumors, respectively; **Suppl. Figure S2D**). Fibroblastic features were seen in *TN* melanomas from UVA-treated animals (3 of 6), but were not overtly apparent in tumors from other *TN* mice (**Suppl. Figure S2E**). Unlike the *TN* model, tumor samples from *TB* mice contained areas of pigmentation, typically characterized by multiple clusters of melanophages distributed at the dermal-hypodermal interface with or without associated neoplastic cells and occasionally within the tumors (**Suppl. Figure S2F**). These results show that although the histopathological features of cutaneous murine and human melanomas differ, a single UVA or UVB exposure can promote the formation of cutaneous, murine tumors with distinct morphologic features.

### Identification of clustered *Flna* and *Map3k1* mutations in *TN* and *TB* melanomas

Prior GEMM studies revealed an enrichment of *Trp53* mutations in melanomas accelerated by full-spectrum-(UVA + UVB) or UVB-irradiation (Viros et al. 2014; Mukhopadhyay et al. 2016;Trucco et al. 2019). However, *Trp53* mutations occur late in human melanoma pathogenesis (Shain et al. 2018). We sought to identify variants associated with earlier stages of melanoma progression, and performed WES using an ensemble calling approach to identify variants in *TN* and *TB* melanomas (see **Methods**). Pooled normals from each inbred mouse model served as germline controls and polymorphisms observed in dbSNP were excluded (Kitts et al. 2013).

Unlike previous reports, *Trp53* mutations were extremely rare in melanomas from our models (1 of 36 tumors, **Suppl. Table 1**). Instead, clustered, recurrent *Flna* and *Map3k1* alterations were observed in tumors from multiple *TN* and *TB* litters (**Figure 2A-C**). Thirteen of the 15 identified *Flna* mutations (87%) localized to the tenth Ig-like repeat of Filamin A (**Figure 2B**). Alterations in this domain are reported to alter Filamin A binding to F-ACTIN and may also affect protein translation and stability (Nakamura et al. 2007; Page et al. 2011; Suphamungmee et al. 2012).

**Figure 2.**
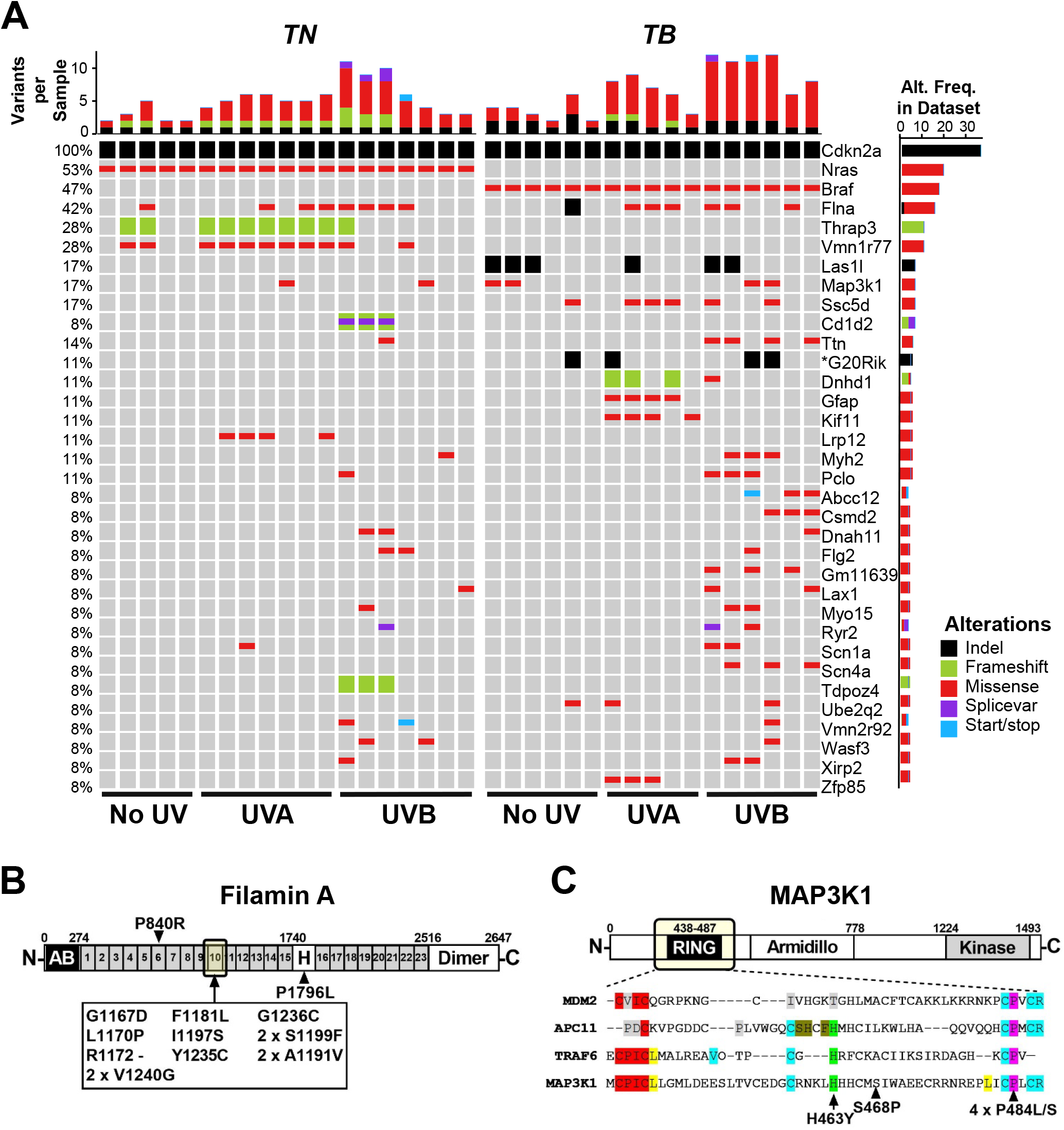
Recurrent genetic alterations in murine models of UV-associated melanomagenesis. **A**, Oncoprint depicting genes mutated in ≥3 *TN* and *TB* tumors treated with no UV, UVA or UVB. Color is used to indicate each mutation type: SNV, indel, frameshift, splice variant or nonsense/stop. Total mutation burden is shown at the top of each sample column. The frequency at which each gene is altered in the dataset is indicated to the right of each row. **B**, Schematic depicting the protein domains of Filamin A. A recurrent cluster of mutations was identified in the tenth Ig-like repeat domain as indicated by the arrow. **C**, Schematic depicting the protein domains of MAP3K1, where recurrent mutations were identified in the RING domain (arrows). Bottom panel depicts a multiple sequence alignment of related E3 ubiquitin ligases, highlighting the conservation of mutated residues.

Two *TN*, and four *TB* melanomas, contained mutations affecting conserved residues of the MAP3K1 RING domain (**Figure 2C**). These findings are consistent with prior publications implicating MAP3K1 and Filamin A in melanoma progression (Ni et al. 2013; Savoy and Ghosh 2013; Mann et al. 2015; Trucco et al. 2020).

### UVB increases the SNV burden of *TN* and *TB* melanomas

In contrast to human melanomas, tumors from genetically engineered mouse models are frequently characterized by a high burden of genomic copy number alterations (CNAs) and few single nucleotide variants (SNVs) (Hodis et al. 2012; Krauthammer et al. 2012; Zhang et al. 2016;Wang et al. 2017; Zloza et al. 2017). Thus, we sought to determine whether a single UVA or UVB exposure caused significant alterations to the genomic landscape of *TN* or *TB* tumors. Fewer CNAs were seen in all *TN* tumors exposed to UVB and in 5 of 7 *TN* tumors exposed to UVA as compared to melanomas from unirradiated controls (**Figure 3A-B**). Conversely, only 3 of 6 melanomas from our UVB-irradiated *TB* mice had a lower CNA burden than tumors from unirradiated controls (**Figure 3A-B**). The most common CNA observed in *TB* tumors was a gain in chromosome 6, the chromosome in which *Braf* resides (**Figure 3A**). Consistent with this observation, 5 of 12 *TB* melanomas showed increased BRAF protein expression as compared to normal, murine skin (**Figure S3**). Recurring copy number gains in chromosomes 1 and 10 were also observed in tumors from the *TB*-UVA, *TN*-No UV and *TN*-UVA groups (**Figure 3A**).

**Figure 3.**
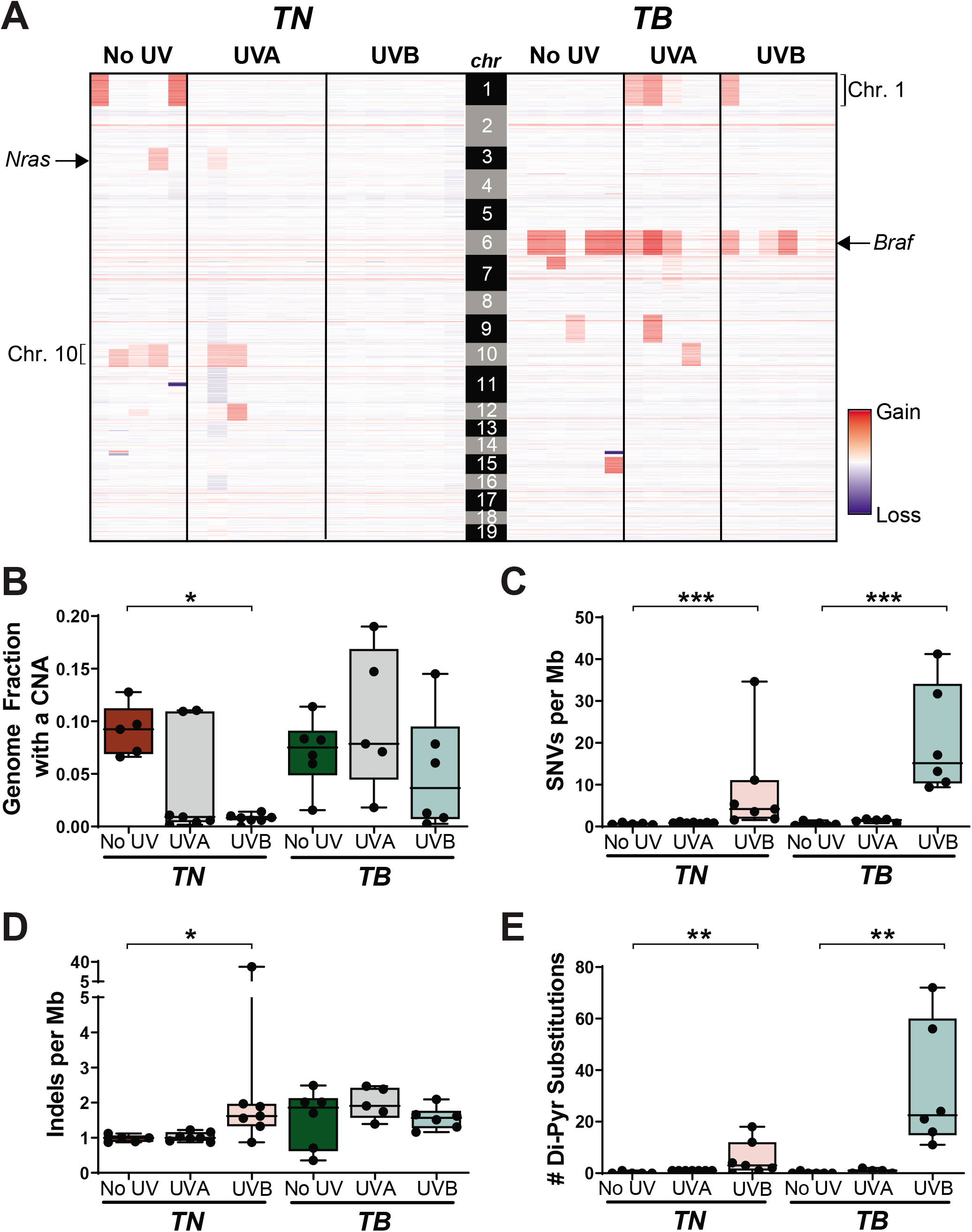
UV alters the genomic landscape of *TN* and *TB* tumors. **A**, Heatmap showing areas of genomic gain or loss within each sequenced tumor. Columns correspond to individual tumors and rows correspond to genomic bins. B, Fraction of each sequenced melanoma genome exhibiting a copy number alteration (CNA), graphed as a box plot with whiskers indicating the 5^th^ and 95^th^ percentiles. Dots represent individual tumors. p-values determined using a Kruskal-Wallis test with Dunn’s correction. **C-D**, Single nucleotide variants (SNVs) (C) and indels (D) per megabase (Mb) of captured genome, plotted and analyzed as in ‘B’. E, Number of dipyrimidine substitutions per tumor, plotted and analyzed as in ‘B’. For **B-E: \***= p<0.05, ** = p<0.01, *** = p<0.001.

The average burden of SNVs increased in both *TN* and *TB* melanomas as a result of prior UVB exposure, whereas the SNV burden of UVA-irradiated *TN* and *TB* tumors was slightly higher, but not statistically different, than that observed in tumors from the No UV groups (**Figure 3C**; 0.67 vs. 0.93 and 1.44 vs. 0.65 SNVs/Mb on average, respectively). The frequency of insertions and deletions (indels) did not differ in tumors from irradiated and unirradiated *TB* mice, but increased in *TN*-UVB melanomas as compared to unirradiated controls (**Figure 3D**). Melanomas from UVB-irradiated *TN* and *TB* mice were also enriched for dypyrimidine substitutions, consistent with the ability of UVB to promote cyclobutane pyrimidine dimer (CPD) formation (**Figure 3E**; (Cadet and Wagner 2013)).

### UVB drives genotype-dependent mutagenesis on the non-transcribed DNA strand

UVA is the most prevalent form of UV in terrestrial sunlight; however, it is poorly absorbed by DNA (Setlow 1974; Sutherland and Griffin 1981; Pfeifer et al. 2005; Khan et al. 2018). By contrast, UVB can directly damage DNA and is the major form of UV responsible for skin erythema and many skin cancers (Setlow 1974). Both bands of the UV spectrum generate reactive species that promote the formation of a wide variety of modified nucleotides (Cadet and Wagner 2013). To examine whether distinct mutation types arise after UVA or UVB irradiation, we quantified the burden of each SNV type (C>A, C>G, C>T, T>A, T>C or T>G) in our sequenced *TN* and *TB* melanomas (**Suppl. Table 2A**). We also examined the prevalence of C>T transitions at CpG sites because methylated cytosines are reported to form cyclobutane pyrimidine dimers (CPDs) with higher efficiency than non-methylated cytosines (Tommasi et al. 1997).

We compared both the absolute number and relative frequency of each mutation type between tumor types from each genotype and UV irradiation status (**Suppl. Table 2B-C)**. As anticipated, melanomas from UVB-treated *TB* mice had a greater number of C>T transitions than UVA or unirradiated controls of the same genotype (**Figure 4A&C**, **Suppl. Table 2B**; *p*<1.02E-14 and 1.82E-13, respectively). The absolute number of C>T mutations was slightly, but not significantly, greater in *TN*-UVB than *TN*-NoUV tumors (**Figure 4A&C**, **Suppl. Table 2B**; *p*=0.22). However, the relative frequency of C>T mutations in both UVB-irradiated models was greater than UVA or unirradiated tumors of the same genotype (**Figure 4B**, **Suppl. Table 2C**; *p*<3.90E-3 for all comparisons). All groups showed a similar number and percentage of C>T alterations at CpG sites, suggesting that methylated cytosines are not preferentially mutated as a result of UVB irradiation (**Figure 4B-C**). Indeed, differences in C>T burden and frequency were primarily driven by mutations at non-CpG sites (**Suppl. Table 2B-C**). The increased frequency of C>T mutations was accompanied by decreases in the frequency of T>C mutations in tumors from both UVB-irradiated models (**Figure 4B; Suppl. Table 2C**). No other mutation types were enriched in a specific genotype or UV irradiation group (**Suppl. Table 2B-C**).

**Figure 4.**
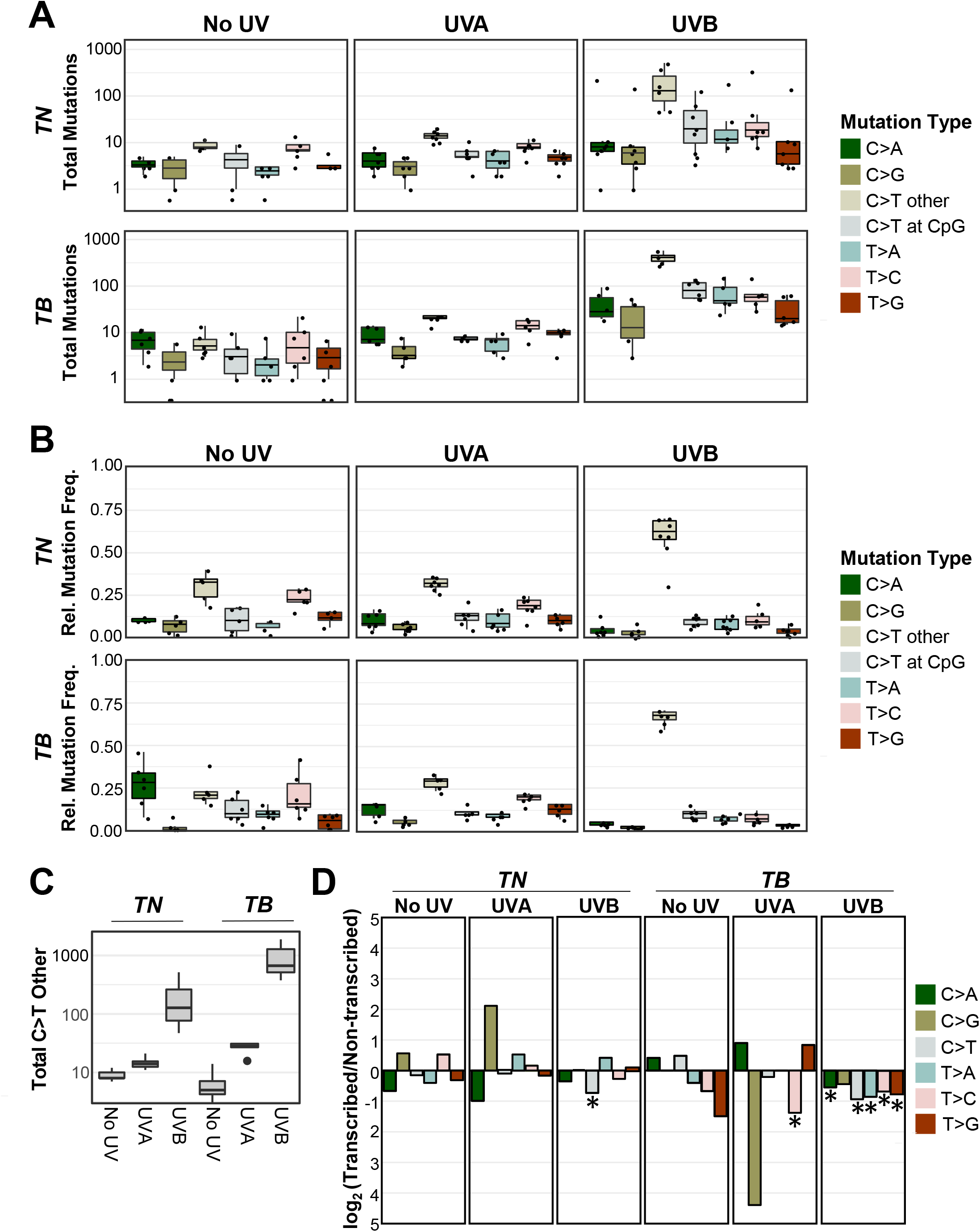
C>T mutations predominate in UVB-induced melanomas. **A-B**, Absolute mutation burden (A) and frequency (B) of each mutation type in *TN* and *TB* tumors arising after mock (No UV), UVA or UVB exposure. See **Suppl. Table 2B-C** for a complete listing of p-values for all comparisons. Statistical significance was evaluated using an ANOVA with Tukey’s HSD post-hoc test. C, Boxplot as depicted in (A), restricted to only C>T SNVs at non-CpG locations. D, The strand location of each mutation type was determined using aggregate data from the indicated mouse models and exposures. Plotted are the log-transformed ratios of transcribed versus non-transcribed mutations. Statistical significance of strand bias was assessed using a Poisson test, where an * indicates significant enrichment. Complete p-value listings are found in **Suppl. Table 2D**.

We looked for evidence of oncogene-dependent mutational enrichments and found that C>T transitions were more abundant (*p*<9.79E-13) in *TB*-UVB tumors than *TN*-UVB tumors (**Figure 4C**, **Suppl. Table 2B**). Other mutation types did not differ significantly in number or frequency between the two genotypes (**Suppl. Table 2B-C**). These data highlight differences in the ability of UVA and UVB to drive melanoma-associated mutations and suggest that an underlying *Braf* mutation may promote the accumulation of C>T transitions.

Studies in cultured fibroblasts and model organisms implicate transcription-coupled repair (TCR) in the rapid repair of UV-induced DNA lesions (Marteijn et al. 2014). Therefore, we investigated whether the SNVs observed in our *TN* and *TB* tumors exhibited a strand bias. Mutations in mock- and UVA-irradiated tumors did not exhibit a strand bias, except in the case of T>C transitions, which were enriched on the non-transcribed strand of *TB*-UVA samples (**Figure 4D**, **Suppl. Table 2D**). Tumors from both UVB-treated models showed a bias for C>T mutations on the non-transcribed strand. C>A, T>A, T>C and T>G mutations were also enriched on the non-transcribed strand of *TB*-UVB, but not *TN*-UVB, tumors. This finding suggests a disparity among *TN* and *TB* melanomas in the biochemistry, incidence or repair of UV-associated DNA lesions.

### Identification of a UVB mutational signature enriched in *TB*-UVB melanomas

CPD-associated C>T lesions are considered classical ‘UVB signature mutations’ and occur preferentially at dipyrimidine sites (Alexandrov et al. 2013a). C>T transitions in other, non-cutaneous cancers lack this specificity (Mitchell et al. 1992). For this reason, we took our sequenced *TN* and *TB* melanomas and quantified the burden of each SNV type within every possible trinucleotide context (**Suppl. Table 3A**). Consistent with these observations, C>T transitions were enriched at TCT and CCT sites in UVB-accelerated *TN* and *TB* melanomas (**Figure 5A & B**, grey bars). UVB also increased the percentage of C>T mutations at other dipyrimidine sites (CCA, CCC, TCA, TCC and TCG) as compared to No UV control tumors in the *TB* model. In contrast to UVB-accelerated melanomas, tumors from mock and UVA-irradiated *TB* and *TN* mice showed a similar distribution of mutations amongst the sixteen potential trinucleotide sites (**Figure 5A & B**). These data are consistent with the pattern of C>T mutations previously observed in a Braf-mutant melanoma mouse model chronically irradiated with UVB (Trucco et al.2019) and prompted us to further explore whether a mutational signature of UVB exposure might be elucidated from our data.

**Figure 5.**
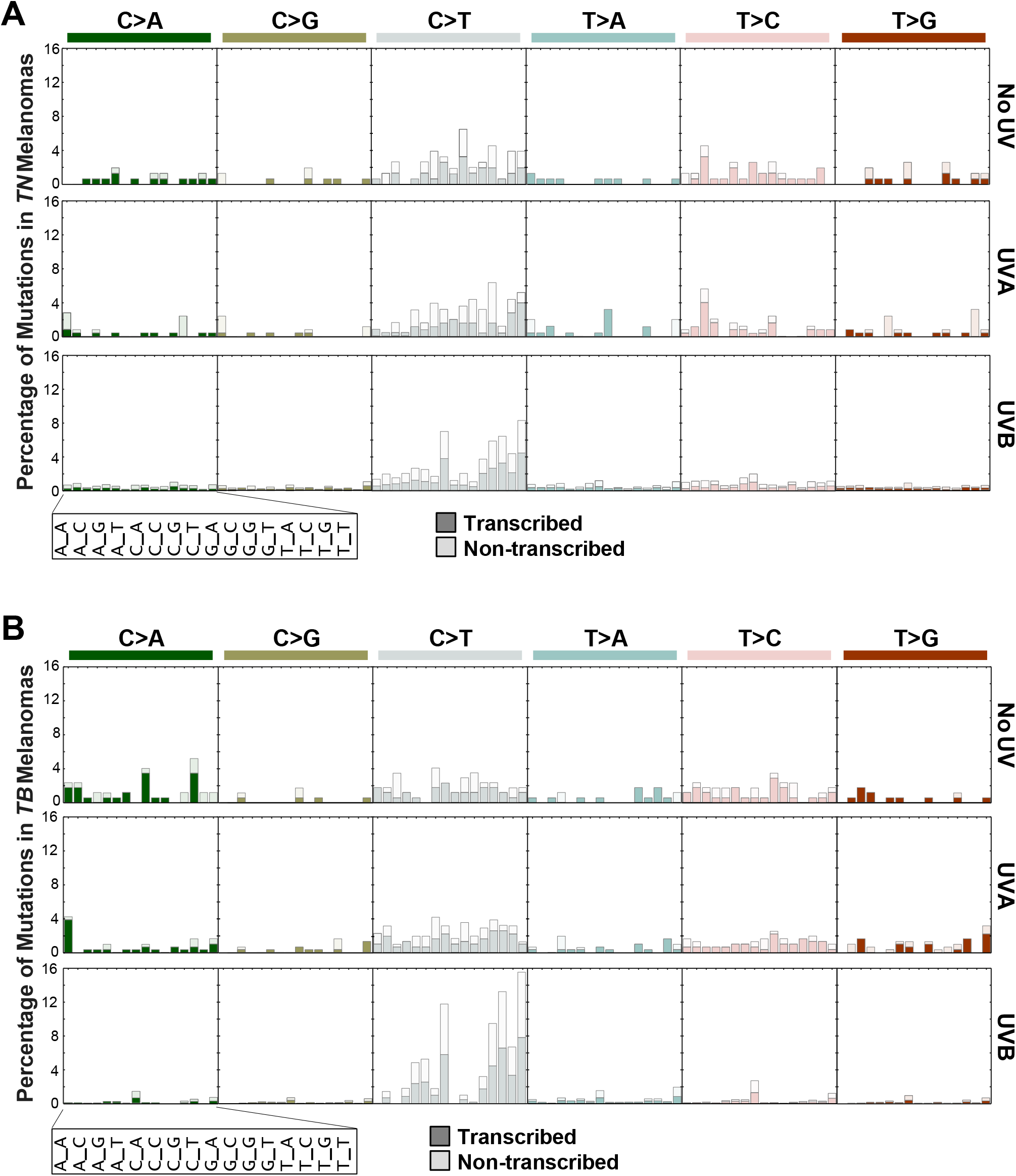
UVB-induced C>T transitions occur within similar trinucleotide contexts in *TN* and *TB* melanomas. **A-B**, Barplots indicating the percentage of each indicated mutation type for a given trinucleotide context in *TN* (A) and *TB* (B) tumors. Each subgraph is a mutation type as indicated, where each column within the graph represents a different trinucleotide context surrounding the SNV of interest. Dark shaded bars represent alterations that occur on the transcribed strand while nonshaded bars indicate alterations that occur on the non-transcribed strand. The height of each bar corresponds to the average number of mutations in the indicated experimental group, normalized to the frequency of the relevant trinucleotide sequence in the mouse exome.

We used SigProfiler (Alexandrov et al. 2013b) to extract co-occurring mutational patterns, ‘mutational signatures’, from our complete tumor dataset. This method consistently identified two distinct mutational processes operational in our *TN* and *TB* melanomas: Signature 1 and Signature 2 (**Figure 6A; Suppl. Table 3B**). The profile of Signature 1 contained an abundance of C>T mutations, with a preference for alterations with a 5’ thymidine (TCT>T<u>C</u>C>T<u>C</u>A>TCG, mutated base is underlined). In contrast, the profile of Signature 2 was relatively flat with no specific mutational preference. To ensure that Signatures 1 and 2 were both reproducible and robust, we used two additional algorithms to extract mutational signatures from our complete dataset: SignatureAnalyzer (Kasar et al. 2015) and Mutational Patterns (Blokzijl et al. 2018). Consistent with SigProfiler, SignatureAnalyzer and MutationalPatterns identified two distinct mutational processes in our dataset (**Suppl. Figure S4**, data not shown). A high degree of similarity was seen between signatures identified by each algorithm, suggesting that the mutational signatures initially found using SigProfiler were robust (**Figure 6B**).

**Figure 6.**
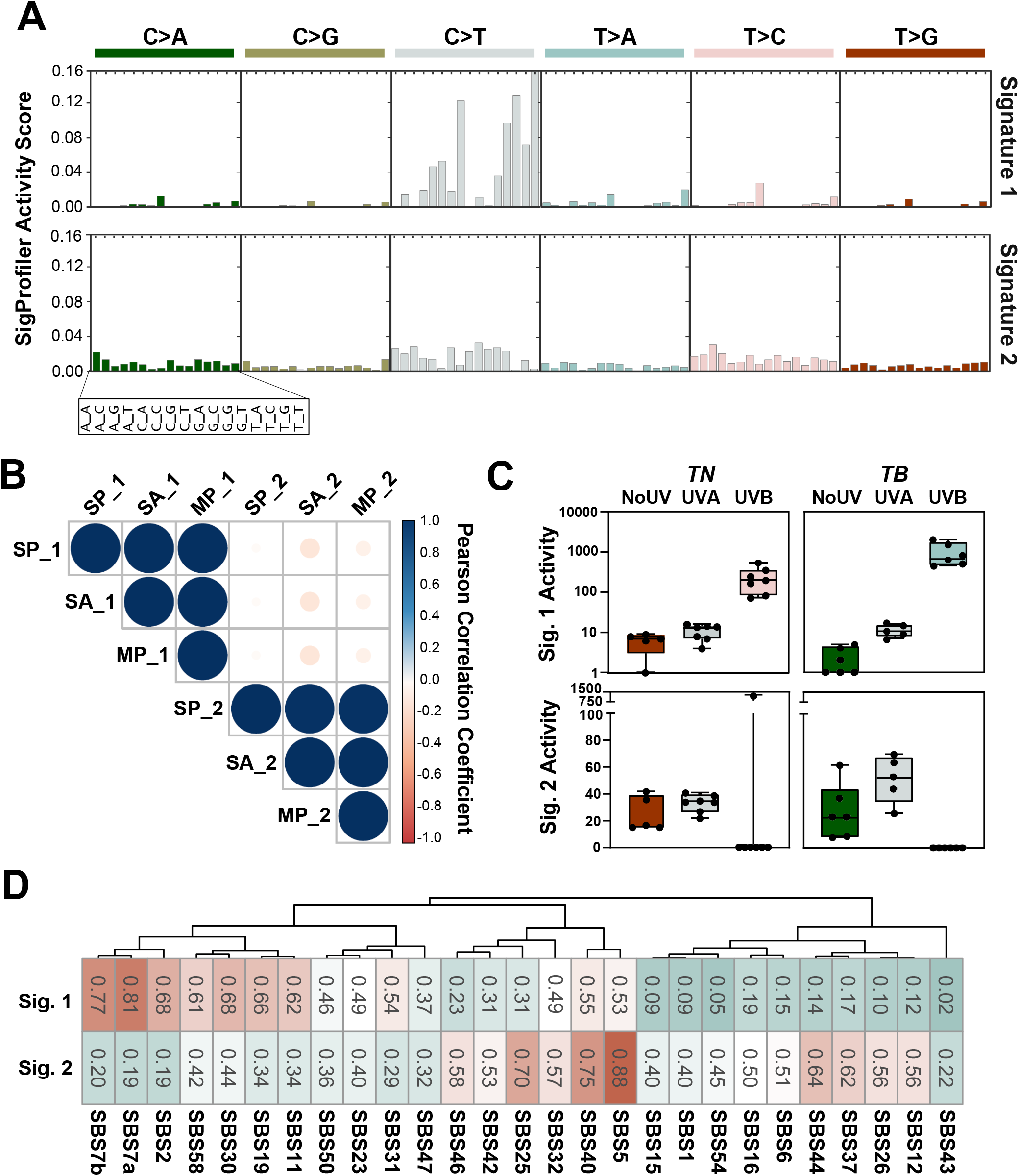
UVB-enriched murine mutational signatures resemble those found in human melanomas. **A**, SigProfiler was used to identify *de novo* mutational signatures in sequenced *TN* and *TB* melanomas. Two signatures were selected based on the average frobenius reconstruction error and signature stability. The y-axis indicates the relative contribution of each trinucleotide mutation type to the discovered mutational signature. Values can be found in **Suppl. Table 3B. B**, Relationships between signatures derived from SigProfiler (SP_1, SP_2), Mutational Patterns (MP_1, MP_2) and SignatureAnalyzer (SA_1, SA_2) are shown. Color indicates the directionality of each correlation with blue indicating concordance and red indicating discordance. The size of each dot represents the absolute value of the correlation. C, Enrichment of SigProfiler signatures in *TN* and *TB* tumors of each exposure type. Significant differences between groups were assessed using an ANOVA with a Fisher’s Least Significant Difference post-test. Complete p-value listings appear in **Suppl. Table 3D** and total mutation counts in **Suppl. Table 3C. D**, Relationship between SigProfiler mutational signatures identified in our dataset (Sig. 1, Sig. 2) and single base substitution (SBS) patterns in the COSMIC database. **Suppl. Table 3E** contains a complete listing of all p-values, which were empirically generated through a cosine similarity permutation test as described in the methods.

We next looked to see if either of the SigProfiler (SP) signatures were enriched in a genotype- or treatment-dependent manner in our mouse melanomas. Signature 2 showed a slight enrichment for pyrimidine transitions, but this enrichement was not specific to any genotype or UV treatment, consistent with the idea that Signature 2 represents background noise or mutagenic process common to all experimental groups (**Figure 6C**). Signature 1 was enriched in melanomas from UVB-irradiated *TN* and *TB* mice over those from No UV- or UVA-treated animals (**Figure 6C**, **Suppl. Table 3C-D**). This enrichment was more pronounced in UVB-*TB* than UVB-*TN* tumors, indicating a greater effect of UVB on the mutational profile of *TB* tumors.

The enrichment of Signature 1 in UVB-treated mice suggested that this profile could exhibit features in common with mutational signatures enriched in sun-exposed human tumors. Therefore, we looked at the cosine of similarity between the signatures identified in our murine dataset and the COSMIC mutational signatures. Our Signature 2 was associated with clock-like signatures that correlate with patient chronologic age, including: SBS40 and SBS5 (**Figure 6D**, **Suppl. Table 3E**; (Alexandrov et al. 2013a; Alexandrov et al. 2015). Our Signature 1 closely correlated with SBS7a and 7b (cosine of similiary 0.81 and 0.77, respectively), which are associated with cancers in sun exposed skin and linked to UV damage (Alexandrov et al. 2013a). Interestingly, there was no significant association between Signature 1 and two other COSMIC signatures, SBS7c and SBS7d, which are enriched in skin cancer and characterized by T>A and T>C SNVs (cosine of similiarity 0.24 and 0.18 respectively) (Alexandrov et al. 2020). These data suggest that mutagenic processes not modeled by our GEMMs may also contribute to human melanomagenesis; however, SBS7a and SBS7b are the predominant signatures found in human melanoma.

## DISCUSSION

Human melanoma has one of the highest mutational burdens of any tumor type (Alexandrov et al. 2013a). Yet, tumors arising in most melanoma GEMMs, including our unirradiated *TN* and *TB* mice, are largely characterized by CNAs rather than SNVs (Hodis et al. 2012; Krauthammer et al. 2012; Zhang et al. 2016; Wang et al. 2017; Zloza et al. 2017). Here we find that a single UVB exposure can resolve this conundrum and effectively recapitulate the high burden of SNVs in human melanoma. Furthermore, the pattern of SNVs in our models is representative of mutational signatures observed in the sun-exposed, human tumors (**Figures 3E, 6C-D**). The UVB-signature derived from our mice does have a higher number of TCT>TTT variants than SBS7a and SBS7b (**Figure 6A**; (Alexandrov et al. 2020)) However, this could be attributed to differences in the trinucleotide frequencies found in each species. Variances in the sequences of transcribed mouse and humans genes could also bias which CPD lesions are efficiently targeted by transcription-coupled repair. Of note, human squamous cell carcinomas deficient in global nucleotide excision repair, exhibit a bias for TCT>TTT variants similar to our models (Chang and Shain 2020). Therefore, differences in how human and mouse cells repair UV lesions may explain the increased prevalence of TCT>TTT variants in our UVB signature. The fact that TCT>TTT variants are enriched in other UVB-accelerated GEMMs further supports this hypothesis (Viros et al. 2014; Mukhopadhyay et al. 2016; Trucco et al. 2019).

Our UVB model recapitulates the burden and distribution of SNVs in human melanoma and identifies recurrently mutated genes seen in the human disease. Specifically, we identify clustered *Map3k1* and *Flna* mutations in *TN* and *TB* melanomas irrespective of UV-irradiation status (**Figure 2**). *Map3k1* is amplified in a subset of human desmoplastic melanomas (Shain et al.2015a)and was previously linked to melanoma progression by two, independent transposon-mediated mutagenesis screens conducted in the Tyr::CreER(T2) *Braf^CA^* model (Ni et al. 2013;Mann et al. 2015). Nevertheless, it remains to be determined how mutations affecting the MAP3K1 RING domain influence tumorigenesis. Similar to MAP3K1, functional defects in Filamin A are implicated in the progression of solid tumors (Savoy and Ghosh 2013). In our models, *Flna* mutations localize primarily to the tenth Ig-like repeat. This domain is responsible for F-actin binding and associated with germline mutations that cause several otopalatodigital spectrum disorders (OSDs) (Moutton et al. 2016). Collectively, these results highlight the potential of forward genetics approaches, like that employed here, to offer insights into the distinct evolutionary trajectories initiated by oncogenic and environmental pressures.

Our results indicate a disparity in the melanomagenic potential of UVA and UVB. Along with prior publications (Noonan et al. 2012; Trucco et al. 2019), these data provide additional evidence that UVA exposures may increase melanoma risk, but not to the same extent as UVB. It is noteworthy that the higher burden of mutations in our UVB tumors did not provide a growth advantage (**Figures 1F, 3C**). Rather, UVB seems to enhance the ability of genetically predisposed melanocytes to initiate tumor formation. This finding aligns with Blum’s interpretation of the kinetics by which UV initiates non-melanoma skin cancers in albino mice (Blum 1969). Specifically, Blum hypothesized that UV-dependent tumor initiation requires a combination of genetic and mitogenic effects. Therefore, in our genetically pre-disposed models, UV-induced growth factors may facilitate initial tumor growth leading to an earlier onset.

What distinguishes this study from past investigations is the direct comparison of UV-carcinogenesis in *NRAS-* and *BRAF*-mutant melanoma. While we did not evaluate the response of healthy tissue, this approach did allow us to detect differences in the mutagenicity of a single UVB exposure among melanomas expressing endogenous levels of mutant NRAS or BRAF (**Figure 2**). It is unlikely that the increased mutational burden of UVB-accelerated *TB* tumors is solely attributed to the longer latency of these tumors as compared to UVB-initiated *TN* melanomas (**Figure 1C-D**). Notably, despite differences in tumor latency, *TN* and *TB* melanomas from the UVA and No UV groups exhibit a similar number of SNVs per Mb (**Figure 3C**). Furthermore, differential enrichment of the UVB mutational signature, as well as dipyrimidine substitutions, suggest that UVB carcinogenesis differs between the two models (**Figures 3E, 6C**).

Whether this difference explains the increased prevalence of *BRAF*-mutant melanomas on parts of the body intermittently exposed to sunlight requires further investigation.

As mentioned in the introduction, a similar proportion of UV signature lesions (C>T and CC>TT) are seen in *NRAS*- and *BRAF*-mutant human melanomas, despite the predominance of *NRAS*-mutant tumors in chronically sun damaged skin (Cancer Genome Atlas 2015). If, as predicted by our data, the presence of a *BRAF* mutation enhances UVB carcinogenesis, then fewer exposures might be required for melanoma progression. Alternatively, *BRAF* and *NRAS-* mutant cells may exhibit differential thresholds for genotoxic stress-induced apoptosis or senescence.

Understanding how *NRAS-* and *BRAF-* mutant melanocytes respond to UVB damage will likely offer insight into the differential SNV burden observed here. Genotype-dependent DNA-damage responses were recently reported in melanoma cell lines (Sauvaigo et al. 2020). However, an earlier report saw no correlation between repair capacity and melanoma genotype (Gaddameedhi et al. 2010). In BRAF-mutant melanomas, loss of p19^ARF^ promotes the epigenetic silencing of *XPC,* leading to deficiencies in nucleotide excision repair (Luo et al. 2013). Meanwhile, pharmacological inhibition of BRAF has been shown to increase nuclear import of the by-pass polymerase, Pol-K, resulting in increased drug tolerance without clear evidence of enhanced mutagenesis (Temprine et al. 2020). Collectively, these results suggest that melanocyte genotype is likely to modulate the cellular response to UV. However, these studies utilized cell lines or extracts, which fail to model the early stages of melanomagenesis and the potential influence of the tumor microenvironment on repair. We posit that isogenically controlled, *in vivo* studies will be necessary to understand how distinct oncogenic and mutagenic stresses cooperate to modulate DNA-damage response and cellular fitness.

## METHODS

### Mouse models

All animal research protocols were approved by The Ohio State University Institutional Animal Care and Use Committee (Protocol #2012A00000134). Mice were backcrossed >7 generations to pigmented, C57BL/6J animals. Inducible knock-in and knockout alleles were activated with 20 mM 4-hydroxytamoxifen on postnatal (p.n.) days one and two as described (Burd et al. 2014). Subjects from each litter were randomly assigned to receive either ambient light (No UV), UVA or UVB on p.n. day three. A single dose of narrow band UVB was delivered to the dorsal side of each animal using a fixed position, 16W, 312nm light source (Spectronics #EB-280C). Based upon the spectrum and intensity of this light source, we calculated the McKinlay-Diffey erythemal effective energy (EEE) of a 4.5 kJ/m^2^ dose, delivered over ~77 seconds, to be 75 mJ/cm^2^ ((Diffey 2002); **Suppl. Figure S5A**). A dose of 75 mJ/cm^2^ EEE UVB is equivalent to 3.9 human minimal erythema doses (MEDs) in an individual with phototype II skin (i.e. someone who tans minimally, but usually burns with red/blond hair and blue/green/hazel eyes) or to approximately 40 minutes of sun exposure when the UV index is Very High [see (Hennessey et al. 2017) for additional information]. UVA was similarly delivered using a 16W source containing two BLE-8T365 bulbs (Spectronics). Based upon the spectrum of these bulbs, the calculated McKinlay-Diffey erythemal effective energy (EEE) of a 70 kJ/m^2^ dose is 14.2 mJ/cm^2^ ((Diffey 2002); **Suppl. Figure S5B**). The average tanning parlor dose is 4.5 Standard Erythema Doses (SEDs; (Dowdy et al. 2011). One SED is equivalent to 10 mJ/cm^2^ EEE-weighted UV light (Diffey 2002). Therefore, an individual receives >3 times more UVA in an average tanning session than a mouse in our 70 kJ/m^2^ experimental protocol (45 mJ/cm^2^ / 14.2 mJ/cm^2^= 3.17).

### Tumor monitoring, processing and histopathology

Protocols for tumor monitoring, processing and immunoblotting appear in the **Supplemental Methods**. Tumor morphology was assessed by a certified member of the American College of Veterinary Pathologists (KMDL) using methods described by Banerjee and Harris (Banerjee and Harris 2000). In each sample the extent of skin and subcutis tumor invasion, tumor pigmentation and the maximum number of mitotic figures were determined from three different fields of view using a 40x objective and 10x ocular lens with a field number of 22 mm.

### Whole exome sequencing

Tumor and germline control DNA was isolated and quality controlled as described in the **Supplemental Methods**. Indexed libraries were generated from 200 ng of genomic DNA using the Kapa Hyper Prep and Agilent SureSelectXT Mouse All Exon target enrichment systems. Exome hybridization was conducted using 500 ng of each DNA library and the resulting target-enriched fragments PCR-amplified (11 cycles). Indexed libraries were pooled and subjected to paired-end 150 bp sequencing on an Illumina HiSeq4000. Average target coverage was 75X (range 53 - 107X). An overview of WES mapping and coverage metrics appears in **Supplemental Table 4**.

### Variant Calling

Sequences were aligned to mm10 using BWA (version 0.7.15) (Li and Durbin 2009). Duplicates were removed using Picard version 2.17.11 and the resulting sequences re-aligned around indels using GATK version 3.6 (McKenna et al. 2010). Variants were called using VarScan2 (version 2.4) (Koboldt et al. 2012), Mutect2 (Cibulskis et al. 2013) and Strelka2 (Kim et al. 2018). Variants identified by all three callers were filtered to remove existing variations in the Ensembl mouse variation database (Yates et al. 2020) and annotated using Variant Effect Predictor (McLaren et al. 2016). Over 200 calls across samples were visually inspected for depth, alignment and read quality in Integrated Genomics Viewer (IGV; (Robinson et al. 2011)). Dipyrimidine mutations were counted as a single event when calculating total mutational burden.

### Analysis of SNVs and CNAs

SNV burden (variants/Mb) was calculated as a function of the total capture region. SNVs occurring within a dipyrimidine sites were counted as a single event. Oncoprints of genes mutated in three of more mouse melanomas were made with ComplexHeatmap version 2.0.0 (Gu et al.2016). To calculate the overlap with human tumors, CNAs were identified using CNVkit (Talevich et al. 2016). Reported CNAs passed a log2 segmentation threshold of 0.2 with support from at least five bins. Genome fraction containing a CNA was determined by computing the footprint of segments surpassing the copy number threshold and dividing this by the total footprint of all segments.

### Mutational spectrum analysis

The total burden and relative contribution of each mutation type to No UV-, UVA- and UVB-induced melanomas was determined using the “mut_type_occurrences” algorithm in the R package for *MutationalPatterns*(Blokzijl et al. 2018). Differences in the absolute number of mutations were assessed using a Mann-Whitney U test. Differences in frequency of each SNV type between UVA or UVB samples versus controls (No UV) were determined using t-tests with Holm’s adjustment for multiple comparisons (p<0.05 considered significant).

A MATLAB implementation of SigProfiler (Alexandrov 2020) and an R implementation of SignatureAnalyzer (Kim et al. 2016) were used to identify *de novo* mutational signatures. Average Frobenius reconstruction error and signature stability were used to select the number of signatures in SigProfiler. The number of signatures selected by SignatureAnalyzer was determined using a Bayesian NMF model described previously (Kim et al. 2016), where two signatures were the most frequent selection from 20 iterations. *MutationalPatterns* was used to examine strand bias and identify *de novo* mutational signatures (Blokzijl et al. 2018). The number of signatures was selected using non-negative matrix factorization, and a rank of 2 was chosen based on maximization of variance explained and cophenetic score. Comparison of *de novo* mutational signatures from SigProfiler and those appearing in COSMIC version 3 (Alexandrov et al. 2020) was completed using a cosine of similarity test, for which empirical *p*-values were generated based on 1,000,000 permutations using the “cosinePerm” function from the PharmacoGx package (Smirnov et al. 2016).

## DATA ACCESS

All raw sequencing data generated in this study have been submitted to NCBI Sequence Read Archive (SRA) (https://www.ncbi.nlm.nih.gov/sra) under accession #PRJNA574176. Code and scripts can be found at https://github.com/bowmanr/UV_mouse_melanoma.

## ACKNOWLEDGMENTS

This work was supported by the Melanoma Research Alliance (309669 to C.E.B.), Damon Runyon Foundation (38-16 to C.E.B; 22-17 to R.L.B.), Pelotonia (R.C.H., E.R.C.) and The National Institutes of Health (F31CA236418 to B.M.M.; R01 R01CA237213 to C.E.B.; P30CA016058 to The Ohio State University).

## Supplemental Materials

**Figure S1.**
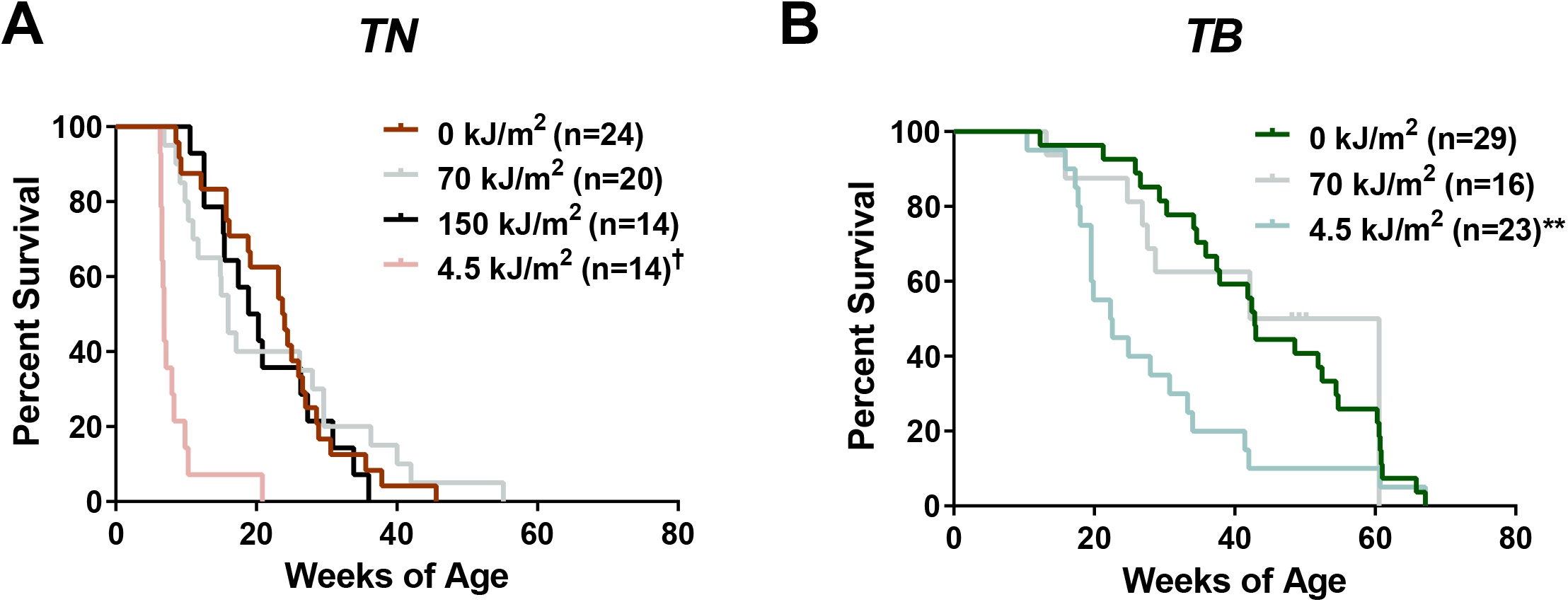
Overall survival of *TN* and *TB* mice treated with No UV, UVA or UVB. **A&B**, Kaplan-Meier curves depicting the overall survival rates of *TN* and *TB* mice treated on postnatal day three with a single dose of ambient light (0 kJ/m^2^), UVA or UVB. Mice were euthanized after meeting pre-determined exclusion criteria due to tumor burden or malaise. Statistical significance was determined by comparing values from control (0 kJ/m^2^) and UV-treated animals of the same genotype using Gehan-Breslow-Wilcoxon tests. † = p<0.0001, ** = p<0.01

**Figure S2.**
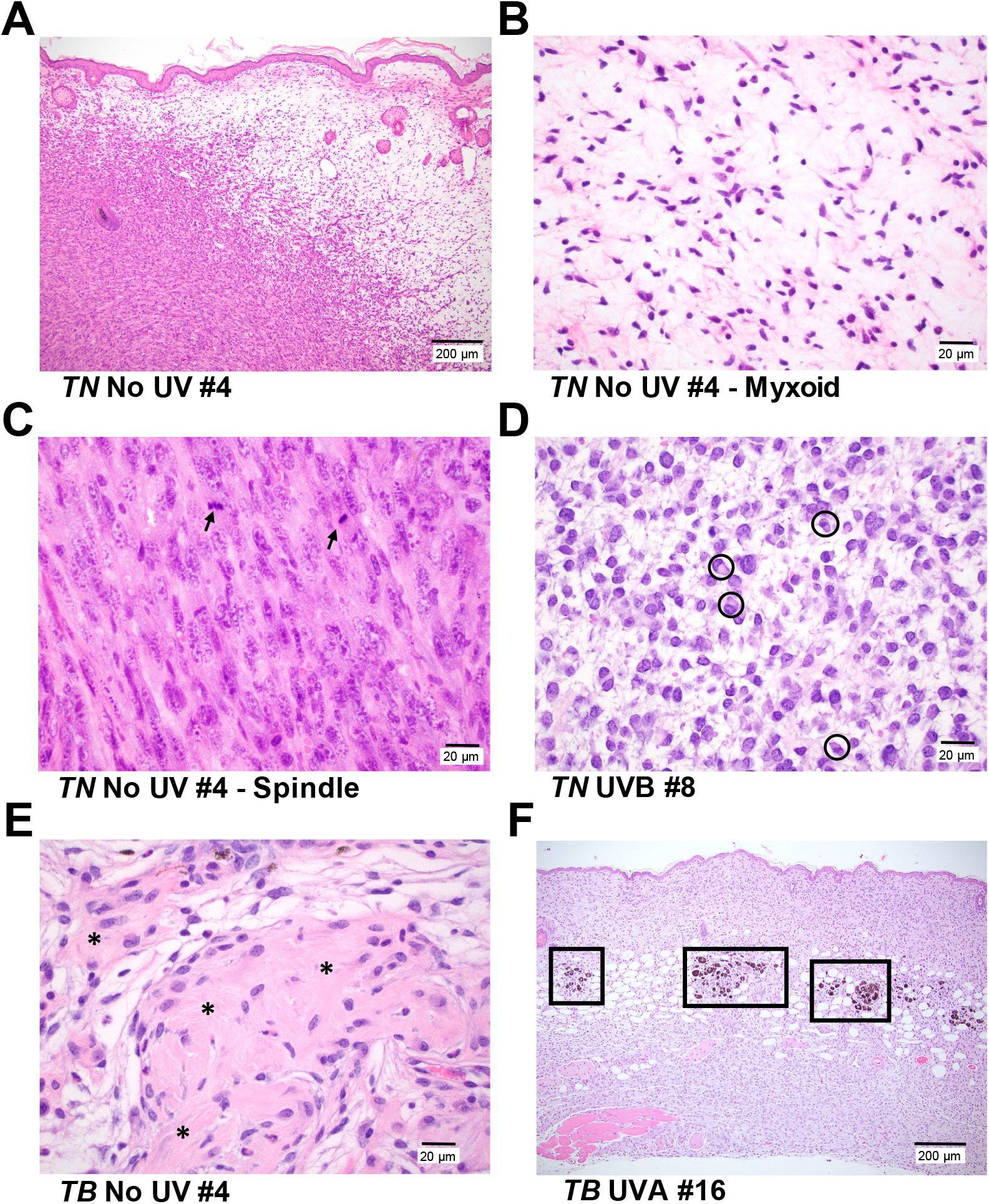
Representative photomicrographs of cutaneous tumors from TN and TB mice. **A**, *TN* and *TB* tumors, regardless of UV exposure history, contain variable proportions of spindle (left, cellular) and myxoid (right, spaces) morphologies. Bar = 200 μm. **B&C**, Higher magnification images of the myxoid (‘B’, bar = 20 μm) and spindle (‘C’, bar = 20 μm) morphologies depicted in ‘A’. Arrows = mitotic figures. **D**, Representative image of the neoplastic cells with plasmacytoid features seen in the majority of *TN* tumors evaluated. (circles; bar = 20 μm). **E**, Half of the tumors in UVA-treated *TN* mice exhibited a fibroblastic phenotype with abundant collagen (indicated by *; bar =2 0 μm). **F**, Tumor samples from *TB* mice contained areas of pigmentation, typically appearing as clusters of melanophages at the dermal-hypodermal interface (boxes, bar = 200 μm). All images are of H&E stained tissues.

**Figure S3.**
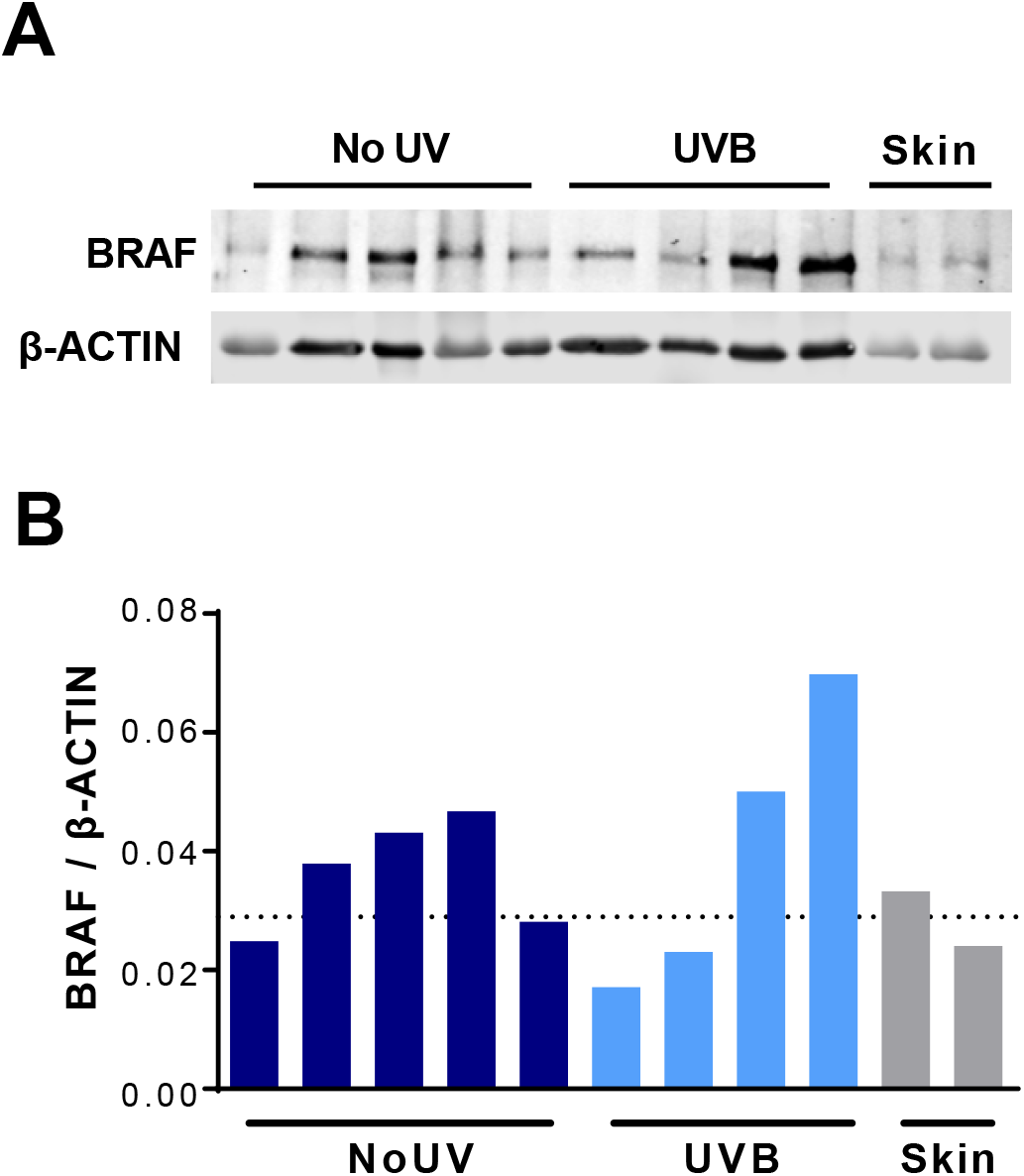
Characterization of BRAF levels in *TB* tumors. **A**, Immunoblot of BRAF and β-ACTIN expression in representative *TB* tumors and whole skin. Each row contains lysate from a single tumor or skin lysate. **B**, Quantification of the immunoblot shown in ‘A’. The dotted line represents the average BRAF expression level in whole skin.

**Figure S4.**
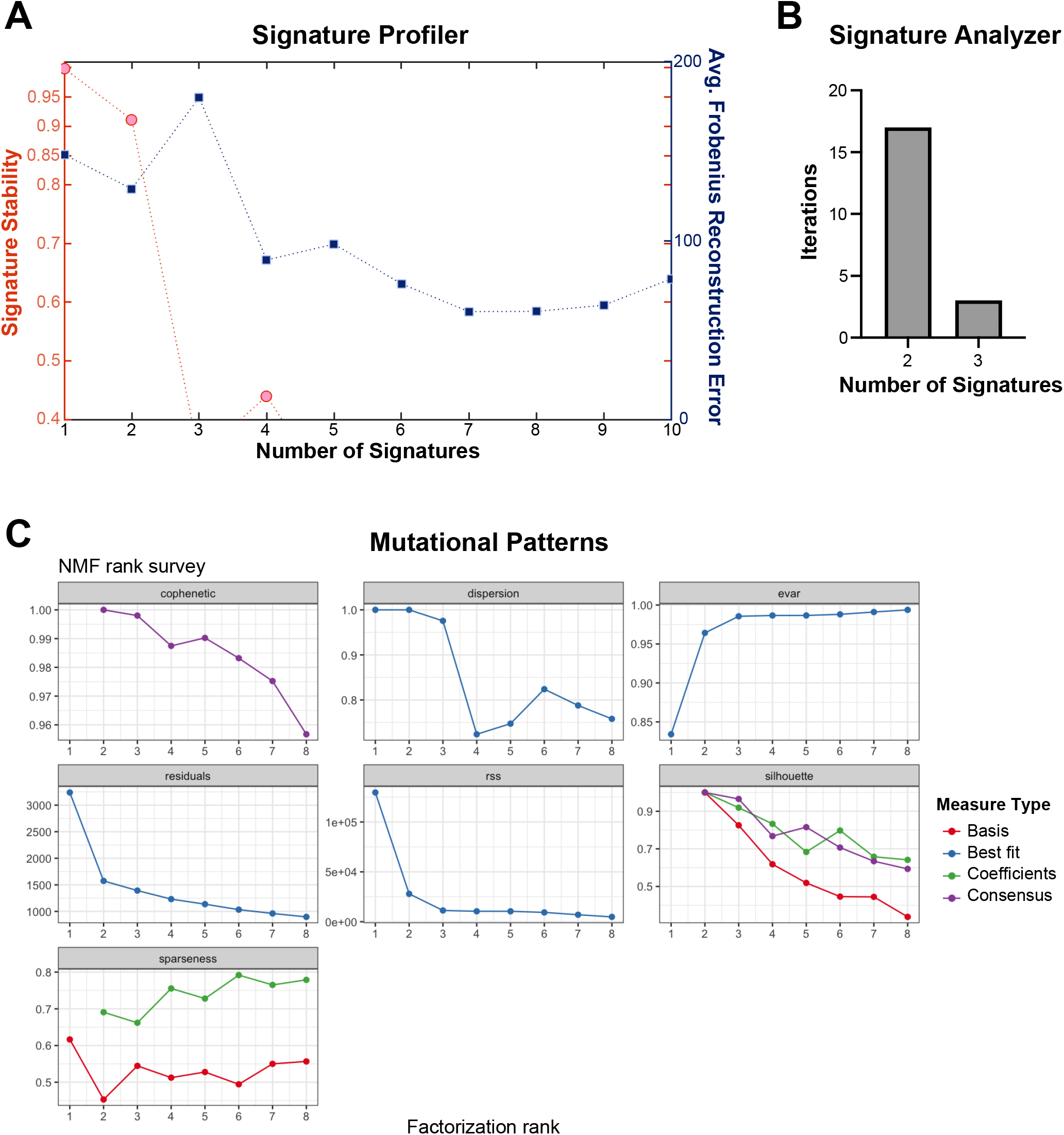
Signature selection metrics. **A**, The mean cosine distance representative of signature stability is plotted on the left y axis, and the average frobenius reconstruction error is plotted on the right y axis for each number of signatures tested by SignatureProfiler indicated on the x axis. **B**, The frequency of signatures identified by Signature Analyzer is shown, with 2 signatures being identified most frequently across the 20 iterations. See methods for details. **C**, Quality metrics produced for NMF estimate as used in MutationalPatterns. Two signatures were selected as the appropriate estimate based on the cophenetic score as described in the methods.

**Figure S5.**
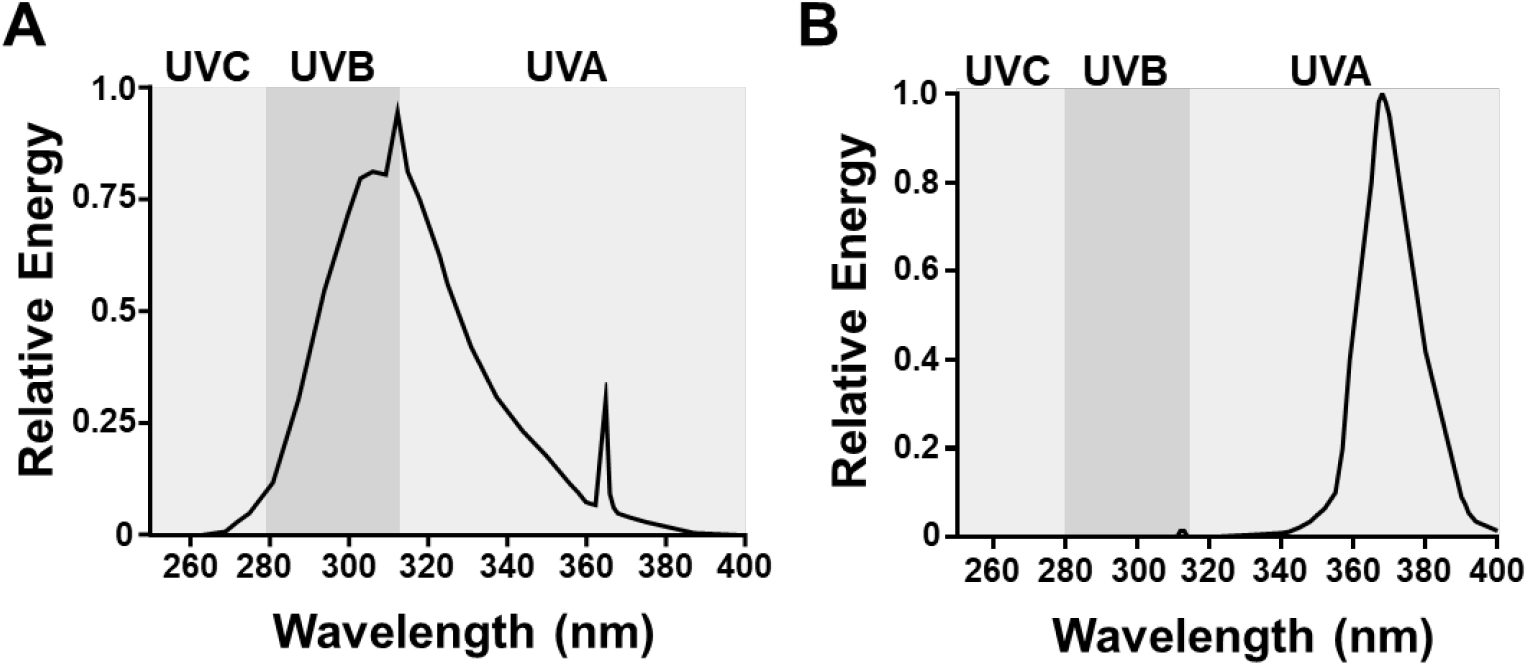
UV light sources. **A**, Shown is the relative amount of energy produced across the UV spectrum by our narrow band UVB, Spectronics EB-280C light source. **B**, Shown is the relative amount of energy produced across the UV spectrum by our narrow band UVAI (340-400 nm), BLE-8T365 light source.

## SUPPLEMENTAL METHODS

### Tumor monitoring and processing

Mice were randomly numbered following treatment and blindly monitored three times a week for tumor formation. Established melanomas were measured by calipers at least three times per week and tumor size (width × length (mm)) recorded until protocol exclusion criteria were met. Representative tumors were harvested from each cohort, fixed in 10% neutral buffered formalin, routinely processed and embedded in paraffin wax. Sections (4 μm) were stained with hematoxylin and eosin (H&E), and evaluated by a veterinary pathologist, certified by the American College of Veterinary Pathologists (KMDL), using an Olympus BX45 light microscope with attached DP25 digital camera (B&B Microscopes Limited, Pittsburgh, PA).

### Tumor immunoblotting

Flash-frozen tumors were homogenized in PBS with Halt phosphatase inhibitor (Thermo Scientific) and protease inhibitor (Sigma) using the Precellys Evolution Homogenizer with Cryolys (Bertin Instruments). The preset elastic setting was used for homogenizing. Tumor homogenates were centrifuged to remove PBS and resuspended in RIPA buffer with phosphatase and protease inhibitors. Lysates were sonicated 2 x 10 seconds using a Branson digital sonifier at 10% amplitude. Samples were centrifuged at 15,000 rpm for 5 min at 4°C and supernatants collected and quantified by Bradford assay (BioRad). Samples (35 μg total) were blotted for BRAF (Santa Cruz sc-5284; 1:500) and β-Actin (Cell Signaling #3700; 1:1000) and imaged using a LI-COR Odyssey CLx system. Bands were quantified using Image Studio Version 5.2 software (LI-COR Biosciences).

### Sample preparation for whole exome sequencing

Tumor DNA was isolated from flash frozen tissue using the Quick-DNA Miniprep Plus Kit (Zymo Research). Tissues were placed in 2 mL tubes containing 190 μL of diluted Zymo Solid Tissue Buffer and 3.0 mm zirconium beads (Sigma Aldrich Cat# Z763902). Samples were then subjected to homogenization using the Precellys Evolution Homogenizer (Bertin Instruments) using the preset elastic setting: speed: 6,800 RPM, cycle: 4 × 30 sec, pause: 45 sec. Homogenized samples were then incubated in 20 mg/mL of Proteinase K overnight at 55°C before continuing with the Solid Tissues protocol as described for the Quick-DNA Miniprep Plus Kit. Control DNA was generated from toe clips or splenic tissue derived from ten representative *TN* and ten representative *TB* animals. These controls were then combined at an equal ratio and concentrated using the Genomic DNA Clean & Concentrator-10 kit (Zymo Research). The integrity and concentration of resulting genomic DNA was confirmed on an Agilent TapeStation.

